# Increasing stomatal density and making the increases broad, near-continuous and quantitative positively regulate *Arabidopsis* growth by utilizing *FSTOMAGEN*

**DOI:** 10.64898/2026.08.12.744549

**Authors:** Yong-yao Zhao

## Abstract

Stomata are the pores on plant surface, and these tiny pores are responsible for the flow of gas between plants and atmosphere. Currently, what effects of the broad and continuous increase in stomatal density achieved via genetic engineering on plant growth and development remain poorly understood.

The 9 *Arabidopsis* transgenic lines with increased stomatal density were acquired through overexpressing *FSTOMAGEN* (the homologs of *STOMAGEN*, which are in *Flaveria*). The intermediate stomatal density (SD) lines exhibited increased trend in biomass. Compared with the lines with low SD, the biomass of *Arabidopsis* lines with intermediate SD (484 mm^-2^) significantly increased. There was a positive and significant correlation between biomass and relative water content. Across these transgenic lines, only during the earlier phase of growth, the leaf area exhibited a gradually increased trend as stomatal density increased, and there was both a significant linear relationship between SD and leaf growth rate and a strong linear relationship between SD and leaf area. In contrast, a clear relationship during the later phase wasn’t observed. Under lower growth light intensity, there was an increased trend of biomass from other lines to the lines with intermediate SD, and the photosynthetic rate and stomatal conductance of the intermediate line were significantly increased.

This study reveals plant-growth alterations that correspond to broad and near-continuous increases in stomatal density achieved via genetic engineering. Our study sheds light on the prerequisites for elevated stomatal density achieved via genetic engineering to promote plant growth.

## Introduction

Stomata are the tiny pores on the surface of plants. They are the passages for the transfer of gas (e.g. CO_2_, H_2_O, O_2_) between plants and atmosphere. A large amount of water lose from plants and spreads into atmosphere through stomata. Therefore, stomata are crucial for controlling the water status of plants. Besides, stomata are also crucial for the assimilation of carbon in plants. Stomata have long been a model to study the development of cells, especially, in plants. The mechanism of stomatal development has been well studied, and many genes related to stomatal development have been found (Pillitteri and Torii 2012; Zoulias et al. 2018). *EPF* family contain a series of genes which encode signal peptides(Takata et al. 2013; Hunt and Gray 2009; Hara et al. 2007; Sugano et al. 2010; Kondo et al. 2010; Hunt et al. 2010). For these genes, the focuses are *EPF1, EPF2* and *STOMAGEN*. The genes in *EPF* family *EPF1/2* negatively regulate the development of stomata, and *EPFL9 (STOMAGEN)* positively regulates stomatal development(Hara et al. 2007; Hunt and Gray 2009; Sugano et al. 2010; Kondo et al. 2010; Hunt et al. 2010). The *EPF1/2* mutant and the overexpression lines have the uniformly changed stomatal density across diverse species(Wang et al. 2016; Hughes et al. 2017; Caine et al. 2018), and *STOMAGEN* mutant and overexpression lines also have the uniformly changed stomatal density across diverse species(Lu et al. 2019)(Tanaka et al. 2013; Shahbaz et al. 2025), suggesting that these genes in *EPF* family are functionally conserved during the evolution.

Adjusting stomatal density could alter stomatal conductance, therefore stomatal conductance changes through genetically manipulating the genes regulating stomatal development(Franks et al. 2015; Tanaka et al. 2013; Jessica Dunn1 2019; Hughes et al. 2017). Consequently, transpiration and photosynthetic rate change (Tanaka et al. 2013; Franks et al. 2015). Furthermore, gradually changed and quantitative stomatal density has been achieved by the genes regulating stomatal development(Karavolias et al. 2024; Karavolias et al. 2023; Sakoda et al. 2020), and lead to specific effects. Recently, a study creates the numerous rice lines with different stomatal density, and most lines have the lower stomatal density than wild type(Karavolias et al. 2024). Stomatal conductance and photosynthetic rate are correlated with stomatal density, indicating that adjusting stomatal density by genetic engineering can quantitatively regulate stomatal conductance and photosynthetic rate(Karavolias et al. 2024). Currently, how increased stomatal density in a broad and continuous manner via genetic engineering affects plant development and growth remains poorly understood.

Currently, increasing plants biomass could capture and save CO_2_ from atmospheric CO_2_ into plants, preventing higher atmospheric CO_2_ concentration and continually increasing atmospheric CO_2_ concentration. Plants biomass is strongly associated with photosynthesis, and increasing photosynthetic rate likely improves plants biomass (Simkin et al. 2015; Kimura et al. 2020). Genetically manipulating stomatal development is an effective way to increase photosynthetic rate(Tanaka et al. 2013; Xia et al. 2022), which probably improves plants biomass(Xia et al. 2022). Indeed, increased photosynthetic rate caused by raised stomatal density leads to increased plants biomass in poplar(Xia et al. 2022). However, the work shows that the overexpression of *STOMAGEN* can’t enhance the biomass of *Arabidopsis*, though stomatal density of the line with overexpression of *STOMAGEN* dramatically increases(Tanaka et al. 2013). One of the possible reason might be the stomata in the transgenic lines are excessive, since it is possible that excessive stomata result in the excessive loss of water in plants and consume the excessive loss of energy which participates in stomatal movement and development. Although there was only one line for each genotype, others work has shown that, the moderately increased stomatal density increases *Arabidopsis* biomass under fluctuating light, the moderately increased stomatal density doesn’t increase biomass under steady light(Sakoda et al. 2020). Although *EPF1* could be used for moderately increasing stomatal density to increase biomass, the function of *STOMAGEN* remains unknown (Sakoda et al. 2020). The changed pattern of the plants biomass where the increases in stomatal density via genetic engineering are broad and continuous remain unknown.

Increased light intensity could increase stomatal conductance. Consequently, although short-term increased light intensity could enhance photosynthesis, long-term increased light intensity might enable plants to experience water stress. Adding further complexity, stomatal density and light intensity exhibit an interplay in their effects on plant growth. Therefore, the growth of the *Arabidopsis* transgenic lines with increased stomatal denisity under different light intensity remains poorly understood. In this study, we examined the growth of multiple *Arabidopsis* (*FSTOMAGEN*) overexpression lines with different stomatal density under different growth light intensity. *FSTOMAGEN* homologs have a quantitative relationship with stomatal density(Zhao et al. 2022), and thus it is chosen for this study.

## Methods

### 1, Building construct, transforming and acquiring the transgenic plants

We overexpressed the functional region of *FSTOMAGEN* (the homologs of *STOMAGEN* in Flaveria) into *Arabidopsis*(Zhao et al. 2022). Specifically, the 35S promoter was used to drive the overexpression of the functional region of *FSTOMAGEN*. The functional region of *FSTOMAGEN* was acquired through the amplification from cDNA of *F.rob*. The signal peptide of the gene in the lines used in this study was also amplified from *F. rob*. These molecular fragments were installed into the pCAMBIA-3300 carrier through homologous recombination(Vazyme). The pCAMBIA-3300 carrier was transformed into E.coli. Then, the carrier was transformed into agrobacterium GV3101(VEIDI). Transgenic lines were acquired by flower-dipping method. The transgenic lines were selected with 2000-diluted Basta. T2 generation of transgenic *Arabidopsis* was performed for subsequent experiment. For the experiment under the lower growth light intensity, the T2 and T3 generation of transgenic *Arabidopsis* were used.

### 2, Growth condition

Two batches of experiments were performed. In the first experiment, *Arabidopsis* were grown in the phytotron (CAS center for excellence in molecular plant science, shanghai, China). The humidity in phytotron was about 60%-65%. The CO_2_ concentration was about 420 ppm. For the experiment, at the beginning, the transgenic *Arabidopsis* lines were grown under about 100 PPFD (μmol・m^-2^・s^-1^). After 19 days, the transgenic *Arabidopsis* lines were transferred into about 300 PPFD(μmol・m^-2^・s^-1^). Then, after 11 days, these transgenic *Arabidopsis* lines were transferred into about 500 PPFD (μmol・ m^-2^・s^-1^). During the whole growth period, sufficient water was given to avoid drought and excessive moistening in soil. In the second experiment, *Arabidopsis* was also grown in the phytotron (CAS center for excellence in molecular plant science, shanghai, China). The humidity and CO_2_ concentration in phytotron were same as the condition in the first experiment. The transgenic *Arabidopsis* lines were grown under PPFD 100 (μmol・m^-2^・s^-1^) at the earlier growth phase, and then after 10 days, the lines were transferred into about PPFD 300 (μmol・m^-2^・s^-1^). During the whole growth period, water was properly given to the *Arabidopsis* lines.

### 3, The measurements of biomass

Biomass was tested by using the weight of Arabidospsis. The weight was tested by two sections: fresh weight and dry weight. The *Arabidopsis* was putted into the oven to be the biomass without water, and the dry weight can be acquired. Both dry weight and fresh weight was measuring by using Electronic Balance (sartorius). The biomass was measured 55days after sowing. We took the biomass above ground.

### 4, The calculation of water content

The water content was calculated with following equation:

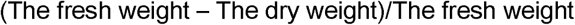

### 5, The photographing of *Arabidopsis*

After transfer the plants to soil, waiting 7 days and then photographing plants, and the interval of the photographing was two days. The photographing was performed by Cannon EOS 1500D camera (Canon Inc., Japan).

### 6, The measurement of photosynthetic gas exchange

The parameters of the photosynthetic gas exchange were measured on the 70th days after the sowing. The Li-6800 (LI-COR) was used to measure the experiments. The fully expanded leaves were used to measure. The measurements were according to the normal instruction. The *Arabidopsis* was adapted to the measument environments for 20 min. The stepwise decreased PPFD (μmol・m^-2^・s^-1^) for A-q: 1200, 1000, 800, 500, 300, 200, 100, 80, 60, 40, 20, 0. Leaf temperature was maintained at 25 ℃. Environment CO_2_ concentration was maintained about 400 PPM. Rapid A–Ci response (RACiR) with continuously ramped reference CO_2_ (Ca) was performed. PPFD was maintained at 600 (μ mol・m^-2^・s^-1^). Leaf temperature was maintain at 24-25 ℃. A-Ci curves were fitted based on Farquhar-von Caemmerer-Berry (FvCB) model. the potential rate of electron transport under saturating light (Jmax) and Maximum velocity of Rubisco for carboxylation were estimated by the fitted FvCB model.

### 7, The observation of stomata

In the second batch of experiment, the mature leaves were used to observe stomata. The leaves were putted on the slides, and then the water was added. The leaves were covered by coverslips, and then the leaves were sent to observe. Since stomata has been observed in previous work(Zhao et al. 2022), we observed one leaf for each line at this time. The stomata were observed by using the optical microscope (Leica, Germany). In the first batch of experiment, the biggest mature leaves were used to observe stomata as the description in (Zhao et al. 2022). We used the artificial intellgence doubao (Bytedance) to measure the stomatal length. 5 stomata along the diagonal were measured, and 16, 11, 11 photos for 3 replicates for 8, 12, 14 line.

### 8, The graphing, plotting, the statistical analysis and linear regression

The one-way ANOVA and Dunnett’s multiple comparisons test were performed by Graphpad prism 8. The student t test was performed by Graphpad prism 8 or excel. The linear regression was performed by Graphpad prism 8 or excel. The growth curve was performed by Graphpad prism 8. The model used in the growth curve was the growth model which is embedded in Graphpad prism 8. The Internal studentized residual and External studentized residual were performed by artificial intellegence, and the artifcial intellegence was name as Doubao which was developed by ByteDance. The graphing and plotting were performed by R language (ggplot) and graphpad prism 8.

## Results

### 1, The transgenic lines with moderately increased stomatal density, which were grown under steady light, exhibited increased rosette area and biomass

The 9 *Arabidopsis* transgenic lines with the overexpression of the gene (*FSTOMAGEN*, the homologs in Flaveria of *STOMAGEN*, Fig 1 C) were acquired. Our work showed that stomatal density in these lines increased stepwise, and there was a positive relationship between stomatal density and expression level(Zhao et al. 2022). Besides, the work from others also indicates that the expression level of *STOMAGEN* among the different lines with the overexpression is positively correlated with stomatal density(Sugano et al. 2010).

**Figure 1.**
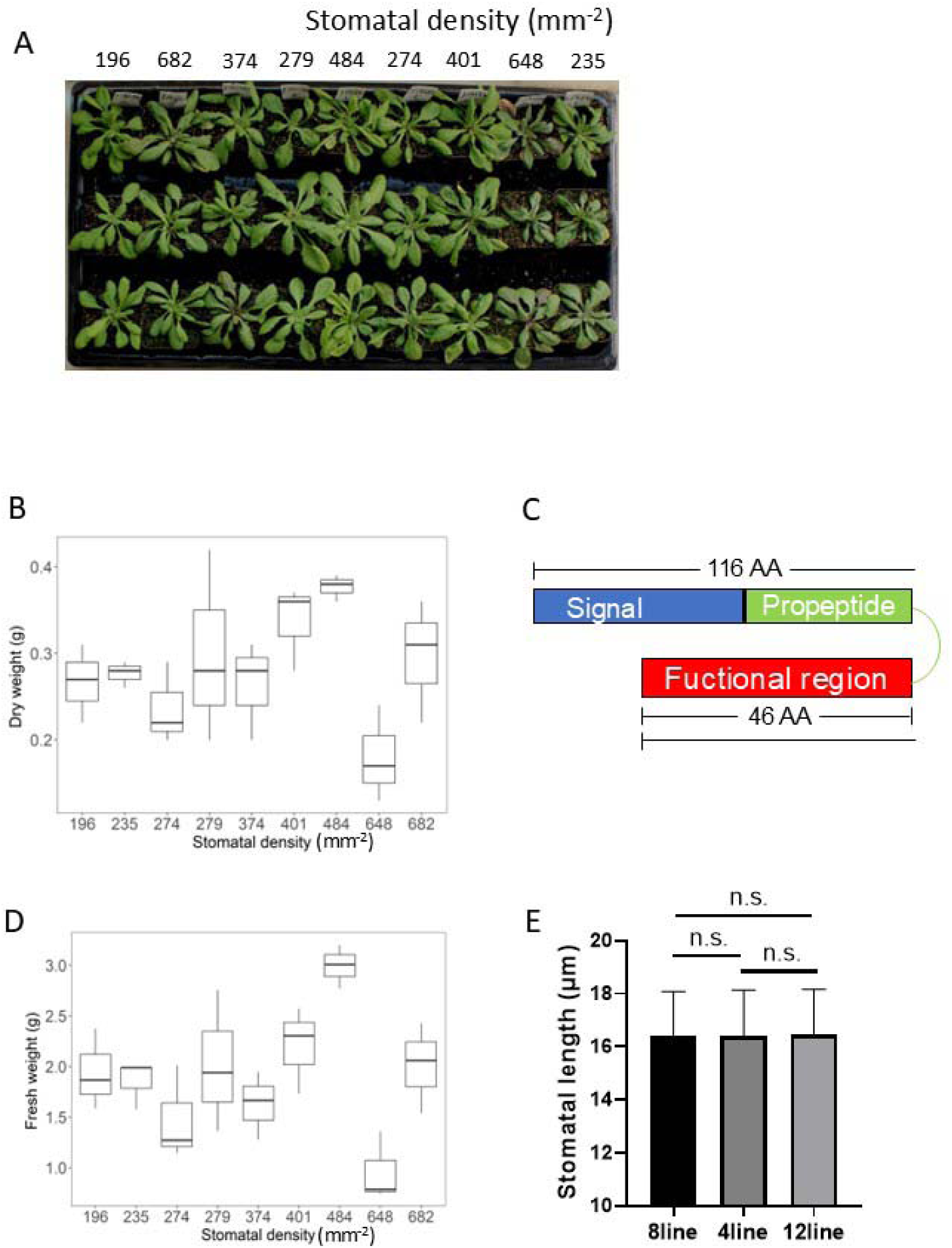
The changes in biomass between different *Arabidopsis* lines with *FSTOMAGEN* overexpression. A, The changes in rosette morphology between different *Arabidopsis* lines with different stomatal density, the stomatal density is shown above the photos of *Arabidopsis*. B, The changes in dry weight between different *Arabidopsis* lines with different stomatal density (n=3). C, The structure of *FSTOMAGEN*, AA indicates amino acid. D, The changes in fresh weight between different *Arabidopsis* lines with different stomatal density (n=3). E, stomatal length of the different *Arabidopsis* lines. The different stomatal density was achieved by the overexpression of *FSTOMAGEN*, and the different stomatal density was achieved by the different expression level of *FSTOMAGEN* in the overexpression. The expression level of *FSTOMAGEN* is shown in (Zhao et al. 2022). The results of the statistical test for Fig B and Fig D are in supplementary file. One-way ANOVA were performed, and Dunnett’s multiple comparisons test were performed for Fig B and D. Student t test (unpaired, two-tail) was performed for the Fig E. * represents P<0.05, and n.s. represents no significant difference in statistics.

Markedly, the differences in the total leaf area between these transgenic lines were observed (Fig 1A). Compared with other lines, two lines (11 and 12 line) had apparently increased total leaf area (Fig 1A). This encouraged us to test the biomass of these transgenic *Arabidopsis* lines. In consistent with total leaf area, there was a variance in biomass between these lines. The trend of the change in fresh and dry biomass, accompanied by the stepwise increase in stomatal density, was first rise and then decrease (Fig 1 B, D). Because the stomatal density of 8 line and 15 line is close to the scope of the stomatal density of wild type(Hronková et al. 2015; Wang et al. 2021), therefore the two lines can be combined as the control. Compared with the combination of 8 line and 15 line, the dry and fresh biomass of 12 line significantly increased (Fig 1B, supplementary file). Compared with the lines with lower stomatal density (8 and 15 line), the medians of fresh and dry biomass in 11 and 12 line were higher (Fig 1 B, D). These results showed that, in order to enhance *Arabidopsis* biomass, there might be a range of stomatal density (401-484 mm^-2^) closing to the thresholds.

We measured the stomatal length of the *Arabidopsis* 8 line, 4 line and 12 line. The stomatal density of 4 line is at the middle between 8 line and 12 line. There was no significant difference in stomatal length between lines (Fig 1E). G_smax_ (maximum stomatal conductance) is determined by the stomatal density and stomatal length. Therefore, the g_smax_ of 12 line and 11 line is significantly and strongly higher than that of 8 line.

### 2. There was a positive correlation between biomass and water content in different transgenic lines of *Arabidopsis*

In the late stage of growth, the rosette of *Arabidopsis* in some lines turned to purple, brown and dried-up (Fig 2A, B). Since these transgenic lines had increased stomatal density, the change in rosette might be caused by the excessive loss of water. Additionally, photosynthesis could be significantly affected by the water content of plants. Therefore, we tested the water content of these transgenic plants. We found that indeed the lines with increased stomatal density had significantly lower water content (Fig 2C, supplementary file). Especially, the water content of the 30 line accompanied with excessive stomatal density was very low (Fig 2C), and the purple and brown of the leaf of this line were most apparent (Fig 2A, B). More important, the leaf of this line became withered (Fig 2A, B). These results indicated that these transgenic lines experienced water limitation during the growth.

**Figure 2.**
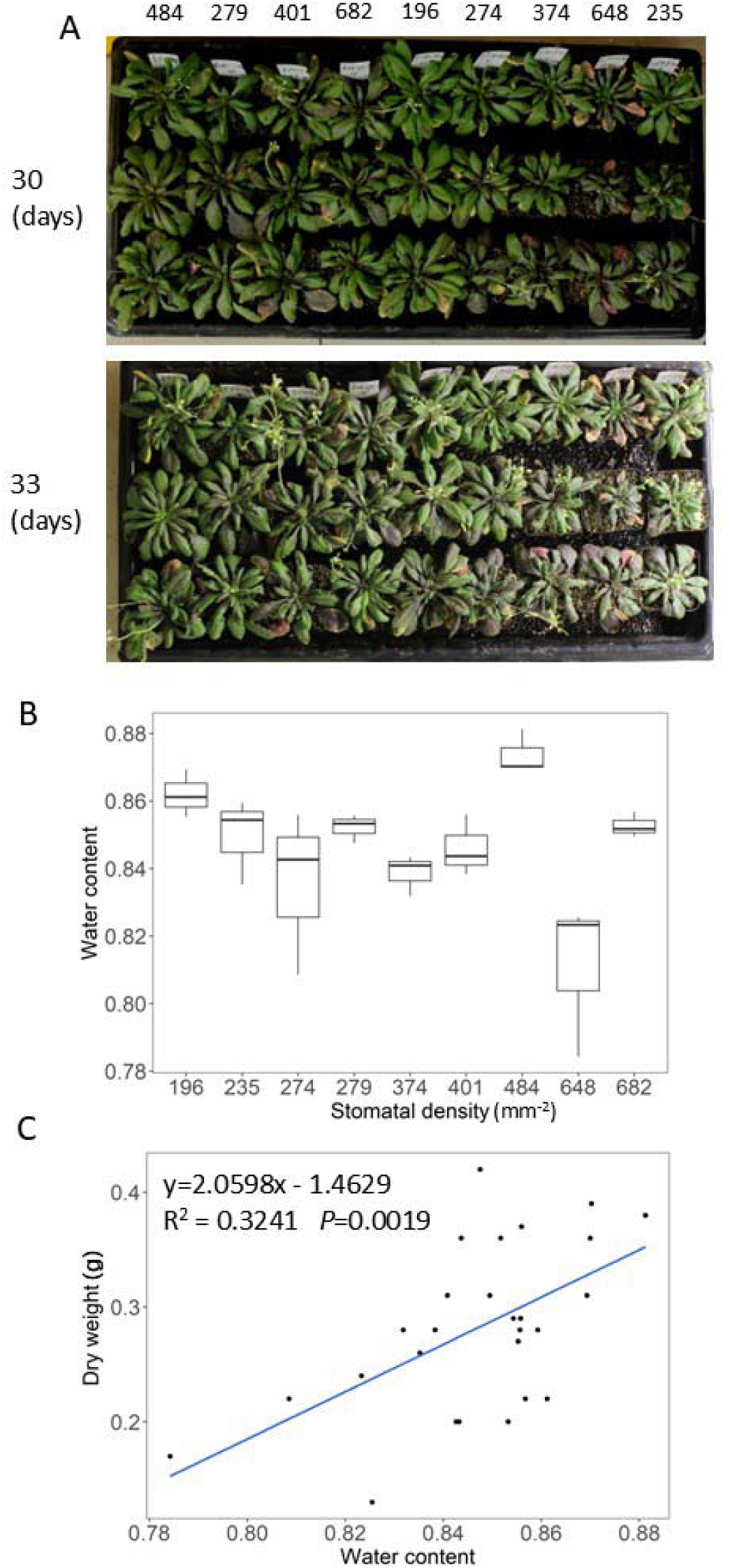
The differences in water content between different *Arabidopsis* lines with *FSTOMAGEN* overexpression. A, The photos for the different *Arabidopsis* lines with different stomatal density at 30 day (above), 33 day (bottom). Stomatal density is shown above the photos. B, The differences in water content between different *Arabidopsis* lines with different stomatal density (n=3). The different stomatal density was achieved by the overexpression of *FSTOMAGEN*. One-way ANOVA were performed, and Dunnett’s multiple comparisons test were performed. The statistical test is in supplementary file. C, The relationship between dry weight and water content is shown.

In the Fig 2B, the lines with increased stomatal density (648 mm^-2^) have the significantly decreased water content as compared with control (Supplmentary file). Then, we performed the sign test. We found the cases that the median of the water content of the lines with increased stomatal density was lower than the median of line 8 occur 7 times, therefore P<0.016, and this result indicated that increased stomatal density reduces the water content of the *Arabidopsis* transgenic lines.

The decreased water content in plants probably decreases the photosynthetic rate, and then decreases the biomass of plants. Therefore, we tested the relation between biomass and the water content of plants. Results showed that there was a positive relationship between dry weight and water content (y=−0.8444+1.2575x, R^2^=0.3038, P=0.0052) (Fig 2D).

### 3. The leaf area and leaf growth rate of the transgenic lines with increased stomatal density exhibited increased trend in the earlier period

There was a large difference in leaf area and leaf growth rate between these lines (Fig 3 A, B, C). The data of the leaf growth was fitted by a growth model, which was approximately an exponential function (Fig 3 D). The leaf growth in the early phase was slower as compared with the leaf growth in later phase (Fig 3 D). However, we didn’t observe a clear relationship between stomatal density and the entire growth (Fig 3A, B, C). We observed the leaf area and leaf growth rate of these transgenic lines during the earlier period. Both the leaf area and leaf growth rate of these transgenic lines exhibited increased trend, accompanied by the increase in stomatal density during this period (Fig 3A, B). This result accorded with the theory that increased stomatal density cause increased photosynthetic rate. It is worth noting that the pattern of the growth rate after 20 days differed from that of the preceding period.

**Figure 3.**
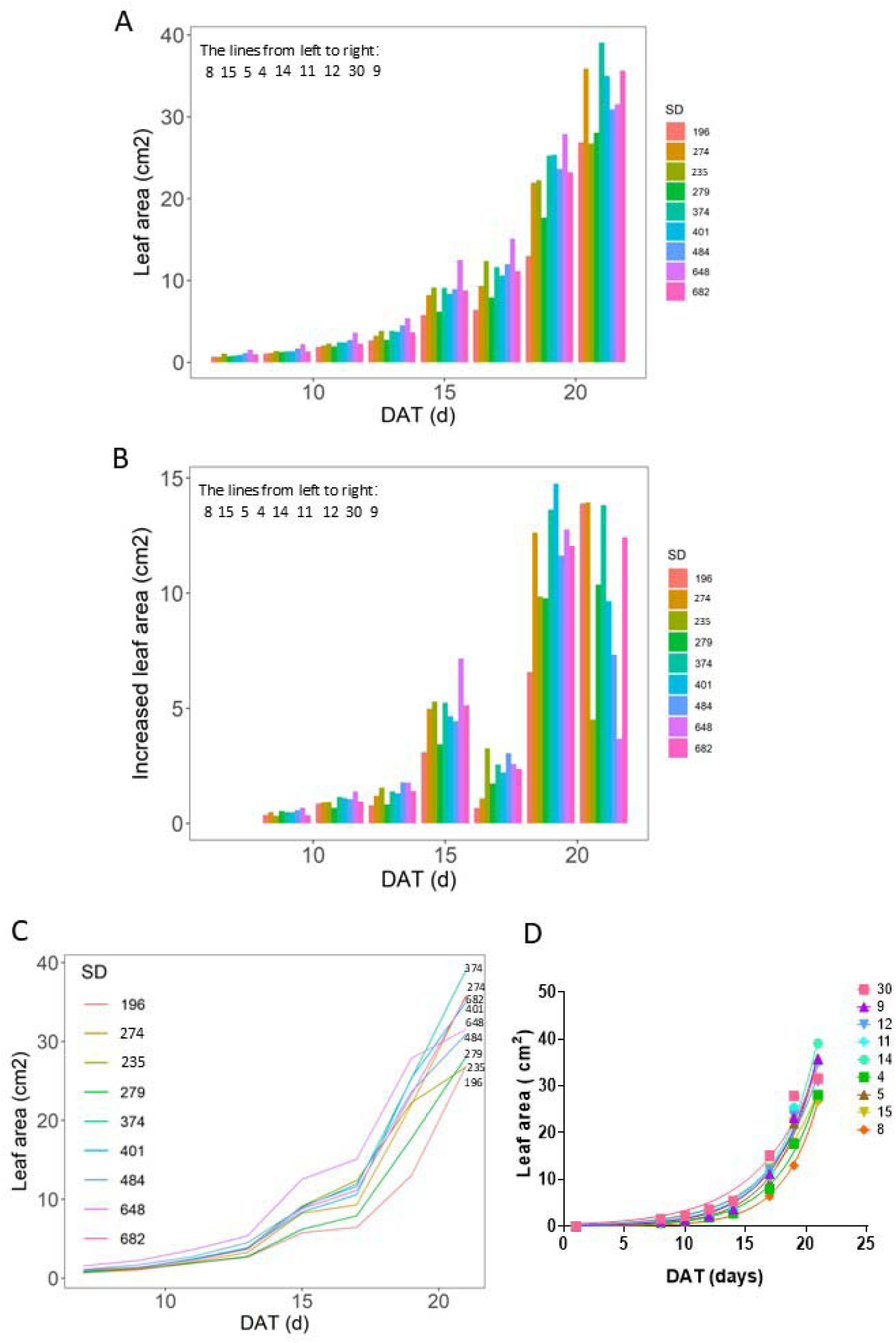
The differences in leaf area between different *Arabidopsis* lines with the *FSTOMAGEN* overexpression at the different date. A, The differences in leaf area between different *Arabidopsis* lines with different stomatal density at the different date. B, The differences in increased leaf area between different *Arabidopsis* lines with different stomatal density at the different date. C, The changed trend in leaf area of different *Arabidopsis* lines during the growth period. The different stomatal density was achieved by the overexpression of *FSTOMAGEN*. D, The fitted growth curve of the growth of leaf area in different transgenic lines. The individual replicates in this graph: 3.

The early phase of leaf growth exhibited apparently linear growth (Fig 5C). There was a significant linear relationship between the slope of the early phase of leaf growth and stomatal density (Fig 5 D). Generally, the leaf growth rate (slope) is positively correlated to photosynthesis, therefore the significant linear relationship between the slope and stomatal density was in line with the notion that increased stomatal density enhances photosynthesis and furthermore biomass accumulated rate (leaf growth rate almost equal biomass accumulated rate), and the increased biomass accumulated rate caused the final difference in the biomass and leaf area (Fig 1, 3), although this case occurred at the early stage (Fig 5D). There was a significantly linear relationships between leaf area and stomatal density at the earlier dates (7-22, 7-24, 7-26 and 7-28) (Fig 4A, B, C, D). We observed an apparent value which likely severally deviated the trend of others (Fig 4A, B, C, D). In order to examine whether it is the real value which severally deviates the trend of others, we performed the analysis of Internal studentized residual and External studentized residual. The results showed that there indeed was a deviated value (Stomatal density is 682 mm^-2^) (Supplementary table 1). Additionally, excessive stomatal density may violate the general rule. Therefore, we have sufficient reasons to dislodge this value from the previous regression. After dislodged this value, it was strikingly that there was a strong linear regression between leaf area and stomatal density (Fig 5A, B), and the R^2^ could reach to about 0.9 (Fig 5 A, B). There was no apparent relationship between stomatal density and the leaf growth in the later phase (Fig 6 A, B).

**Figure 4.**
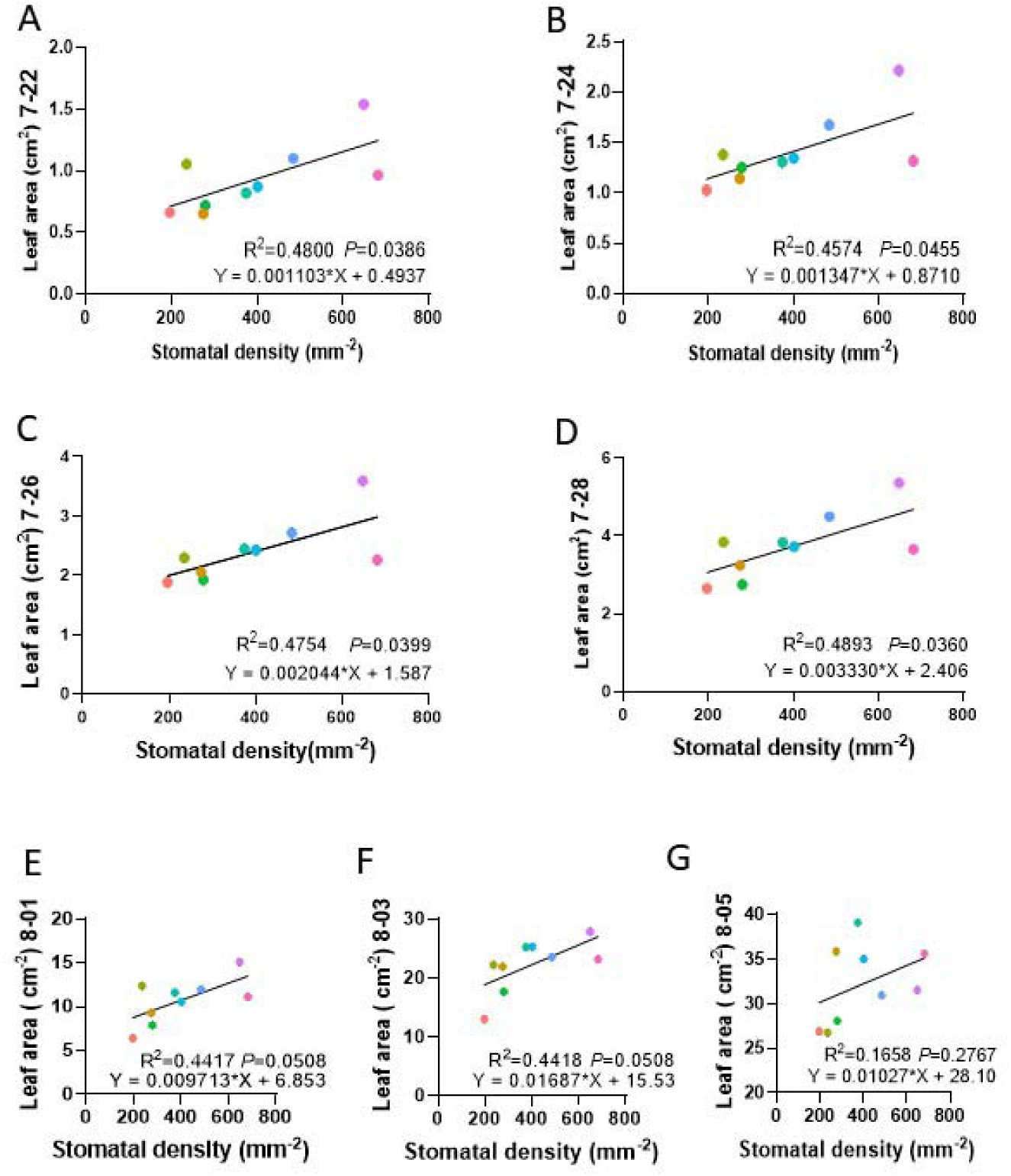
The relationship between leaf area and stomatal density. The relationship between leaf area and stomatal density at 2022-7-22 (A), 2022-7-24(B), 2022-7-26(C), 2022-7-28(D), 2022-8-01(E), 2022-8-03(F), 2022-8-05(G). The regression function, P value and R^2^ were within each panel. The individual repicates: 3.

**Figure 5.**
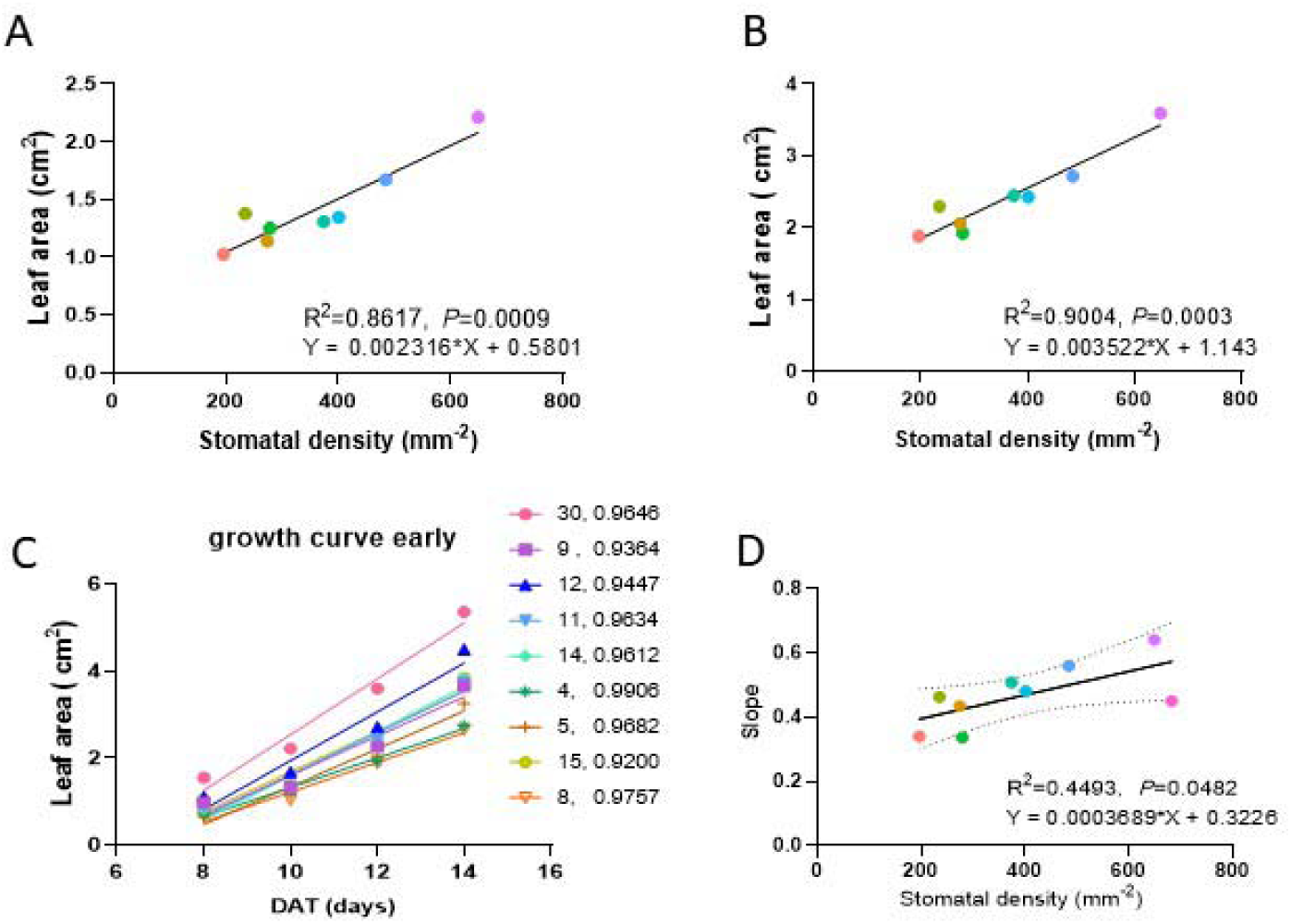
The differences in leaf area between different *Arabidopsis* lines during the early growth phase. A, The relationship between stomatal density and leaf area at date 7-24 without the 682 mm^-2^ stomatal density. B, The relationship between stomatal density and leaf area at date 7-26 without the 682 mm^-2^ stomatal density. C, The linear regression between the days and leaf area in the early phase of growth for the different lines. The number of lines and R^2^ were shown in the right top corner in this panel. D, The relationship between stomatal density and slope (leaf growth rate in the early phase). The individual replicates in this graph: 3.

**Figure 6.**
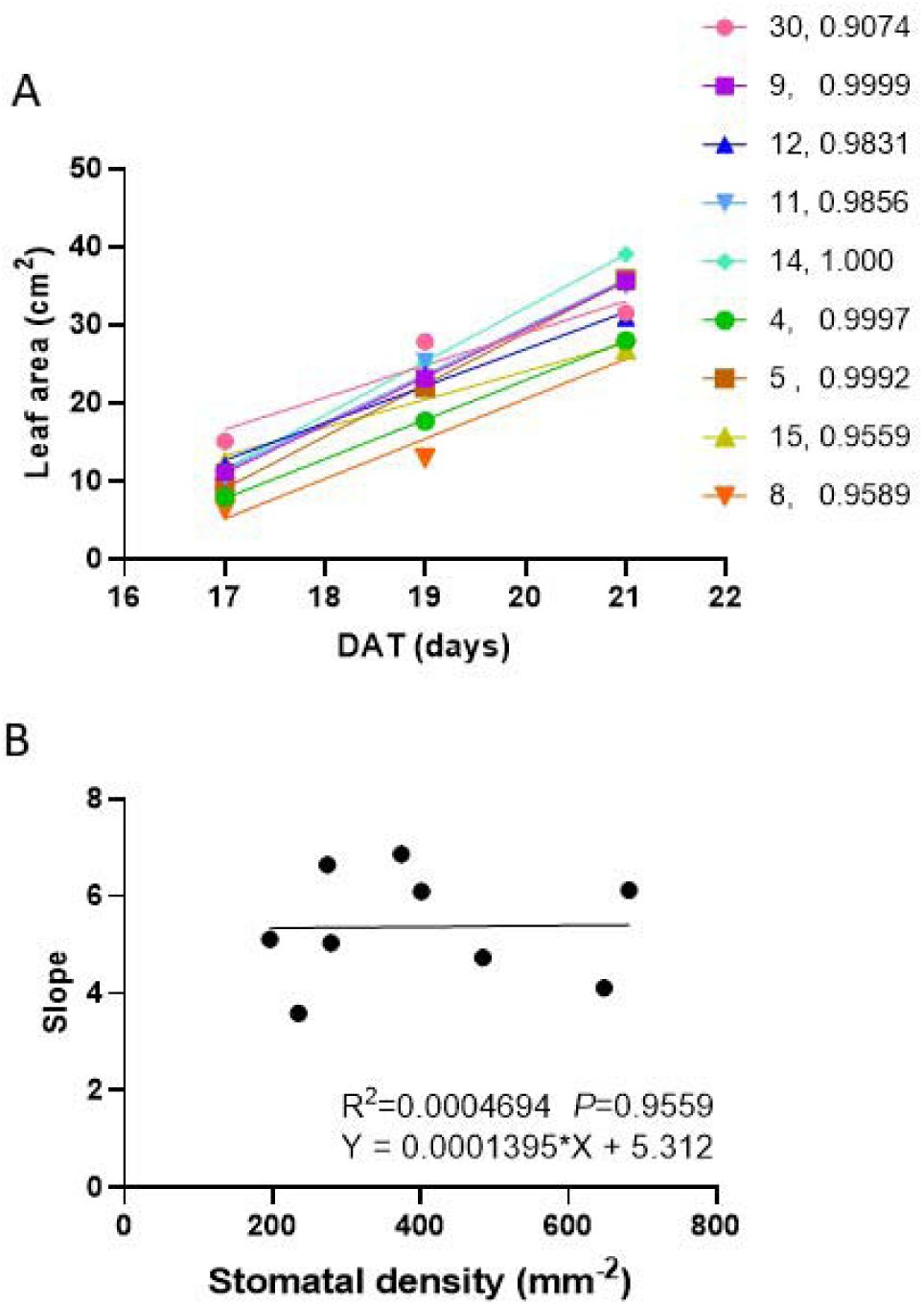
The leaf growth rate in the later growth phase A, The linear regression between the days and leaf area in the later phase of the growth for the different lines. The number of lines and R^2^ were shown in the right top corner in this panel. B, The relationship between leaf growth rate (slope) and stomatal density. The individual replicates: 3.

### 4, The differences between the transgenic lines were less when lines grew under the lower light intensity

The transgenic lines were grown again to explore the phenotype under other condition, and the light intensity was adjusted to lower level (100 to 300 μmol・m^-2^・s^-1^). We found that there are differences in plants biomass and photosynthetic rate between the certain transgenic lines in this experiment (Fig 7 A, supplementary fig 5 A, B). The median of line 14 was higher than that of line 8 (Fig 7 A, D). The difference between line 14 and line 8 was at the statistic significant margin (P=0.08), which was likely caused by the large variance (Fig 7 D). Nonetheless, the higher median of line 14 also accorded with the notion that moderately increased stomatal density enhanced plants biomass. The biomass of 9 line was significantly lower than 8 line, and there was no significant difference between biomass of 30 line and 8 line (Fig 7 A, D). The stomatal density of certain plant in 30 line in this experiment was found slightly increased (Supplementary figure 5), which was probably caused by the gene silence, and the stomatal conductance exhibited significantly increased in the plants of 30 line (Supplementary fig 6 B). The photosynthetic rate of 14 line was significantly higher than that of 8 line, and the stomatal conductance exhibited the identical pattern (Fig 7 C, E, F). There was no significant difference in Vcmax and Jmax between 14 line and 8 line. (Fig 7 B).

**Figure 7.**
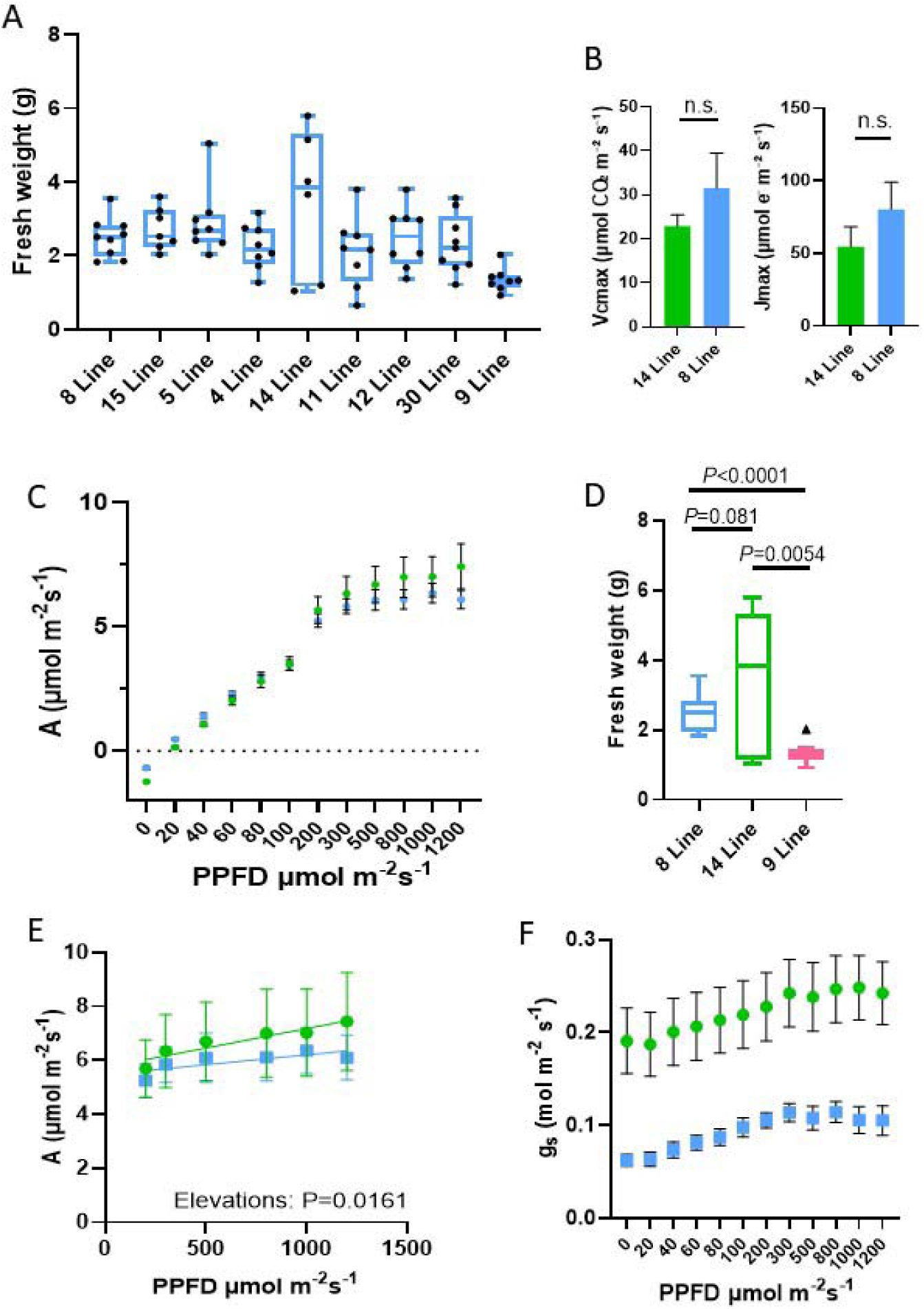
The biomass of the *Arabidopsis* transgenic lines under the lower light intensity (300μmol・m^-2^・s^-1^) A, The fresh biomass of the *Arabidopsis* transgenic lines under the lower light intensity (300 μmol・m^-2^・s^-1^). n≥6. B, The Vcmax and Jmax (14 line and 8 line). n=3. C, A-q curve (14 line and 8 line). n≥4. D, The fresh biomass of the 14 line and 8 line. n≥6. E, The later phase of the A-q curve (14 line and 8 line). n≥4. F, Stomatal conductance (g_s_) of the 14 line and 8 line under different light intensity. n≥4. Student’s t-test was performed. n.s. denotes no significant difference.

## Discussion

### 1) Promoting stomatal development to 8 different levels by overexpressing *FSTOMAGEN* could positively regulate *Arabidopsis* development and growth

In this study, we focused on the broad, near-continuous and quantitative increase in stomatal density by genetically manipulating the expression of the gene (the homolog of *STOMAGEN*, the homolog is from Flaveria genus). Although the increased stomatal density could short-term increases the photosynthetic rate (by increasing C_i_), it is more probable that the increased stomatal density causes the long-term water stress(Doheny-Adams et al. 2012), and the water stress reduces long-term photosynthetic rate, and thus reduces the biomass(Tulva et al. 2024; Tanaka et al. 2013; Doheny-Adams et al. 2012). Our results showed that the lines with moderately increased stomatal density (484 mm^-2^) had significantly higher biomass as compared to the lines with lower stomatal density (Fig 1 B, D). The increased stomatal density in our work caused increased g_smax_(Fig 1), and our growth condition ensured the stomatal opening as much as possible (high light, suitable water condition), therefore the increased g_smax_ should increase the photosynthetic rate and further biomass (Fig 1B, D). It exhibited the increased trend in biomass of the intermediate stomatal density lines (401 and 484). Therefore there might be the thresholds of stomatal density (closing to 401-484 mm^-2^). That the biomass is not increased by mildly increasing stomatal density is likely attributed to the following reasons(Sakoda et al. 2020): the increased stomatal density doesn’t reach the thresholds; the used gene differs (*FSTOMAGEN* is different from *EPF1*).

We found that, during the earlier phase of growth, the leaf area and leaf growth rate of *Arabidopsis* transgenic lines exhibited a increased trends as stomatal density increased (Fig 3A, B Fig 4 and Fig 5), but the changed trend during the late phase is ambiguous (Fig 6). Further, there was a strong and significant linear relationship between leaf area (or the slope of the leaf area growth) and stomatal density in the earlier phase of *Arabidopsis* growth (Fig 4, 5), suggesting that, compared to the other lines, the final increased biomass of intermediate lines was caused by the increased stomatal density. Under the normal growth conditions, there is a strong correlation between leaf area and biomass, therefore our studies suggested that the overexpressing FTOMAGEN could quantitatively and positively regulate the biomass of *Arabidopsis* during the earlier growth phase. Ultimately, our study suggests that prerequisites, e.g., magnitude of increase or growth stage, are essential for promoting plant growth under genetically elevated stomatal density. This is also reflected by the results of different light intensities (Fig 1 and 7).

In agreement with other work(Tanaka et al. 2013), our work also found that biomass remained unchanged or reduced in the lines with drastically increased stomatal density (Fig 1 C, D, supplementary fig 7 D). The reason might be that the excessive stomata strongly enhance transpiration (Fig 2C), and water loss through transpiration largely exceeded water uptake through root. Consequently, the plants were subjected to a severe water deficit (Fig 2 C). The severe water deficit would cause two occasions: 1, inducing the decrease in stomatal conductance; 2, weakening the biochemical reaction in photosynthesis. Consequently, the growth of the plants was hindered and the biomass was reduced. The notion that the decreased biomass was caused by the decreased water content in this study was supported by (Fig 2D). Another reason for the decreased biomass of the lines with extremely increased stomatal density should be that the energy was overly consumed in the stomatal development and movement. The excessive energy consumption reduced the biomass which should have increased in the lines with excessive stomata.

### 2) Genetically manipulating *FSTOMAGEN* promisingly benefits the reduction of the concentration of atmospheric CO_2_ and production

The genes in regulating stomatal development are functionally conserved across different species (Yin et al. 2017; Lu et al. 2019; Wang et al. 2016; Caine et al. 2018; Hughes et al. 2017). This means that we can adjust stomatal density in various species through genetically manipulating the homologs of the genes. Additionally, our work has shown that the substitution of signal peptide does not affect the function that regulates stomatal density(Zhao et al. 2022), indicating the function of the functional region is independent from the function of the signal peptide. Therefore, various signal peptides that can accurately locate the gene may be able to replace the own signal peptide of *STOMAGEN* homologs.

The increase in the concentration of atmospheric CO_2_ accompany the raised atmospheric temperature (Tonn 2007). The accelerated increase in the concentration of atmospheric CO_2_ could accelerate the change in global climate and environment, causing the extreme and rare climate and environment (Including the natural hazard like drought and flood). Therefore, it is necessary to explore the approach to prevent the increase in the concentration of atmospheric CO_2_. Our study indicated that, given the required prerequisite, increased stomatal density promote *Arabidopsis* growth and improve the biomass of *Arabidopsis* (Fig 1 B, C, D), suggesting that increased stomatal density is a promisingly way to enhance plant’s capacity to sequester external CO_2_. Because the genes regulating stomatal density are functional conserved among various species, it is probably feasible that expand this work to various species, and then benefit the decrease in the concentration of atmospheric CO_2_. Especially, a large number of species have C_4_ photosynthesis. It is very attractive for us to test C_4_ plants, since C_4_ plants intrinsically already have strong growth and high production of biomass. The increased biomass of plants can be useful for human. For example, in the industrial and manufacturing sectors, the increased biomass can be used to produce some productions that need cellulose.

### 3) An example for the cooperation between genetic engineering and the controlling of ambient environment

The traits enable plants to fit certain ambient environment, and this is done by evolution. But currently, the changes in geographical environment become faster. A faster strategy for the adaptation is required to meet the faster changes, which could be realized by genetic engineering. Manipulating *FSTOMAGEN* enables plants to fit the geographical environment with sufficient water and high light intensity, enhancing quick fitting. Additionally, the Intelligent plant factory is another strategy to meet the fast changes in geographical environments. In future, the environments (sufficient water and high light intensity) in the intelligent plant factory will be suitable to the increased stomatal density.

In theory, the characteristics are more complex (For example, biomass), it will be more difficult to enable plants to have the expected characteristics through using single gene. Additionally, most current goal of the genetic engineering is to find the genes that function in most and even all environments, which seems not realistic. This largely restricts the application and the development of genetic engineering and molecular biology. It is believed that most genes have their roles, and when in the corresponding environment, they can exhibit their functions in regulating the traits. Our study proposed that it is an effective solution that the cooperation between the genetic manipulation and the controlling of ambient environment is conducted. In this study, the increased extent of the biomass of *Arabidopsis* was higher under higher light intensity and sufficient water (Fig 1, Fig 7). Lower light intensity could weaken biomass improved by increased stomatal density since the photosynthetic biochemistry is limited. Growth condition (high light intensity and sufficient water) is crucial for the increased biomass accompanied by increased stomatal density (Fig 1, fig 7 A). Therefore, this study is a valuable example and provides the new insights for these kinds of studies.

### 4) Stomatal apparatus is a model for the age of quantitative biology

Genetic engineering in biological science has long been a research hotspot. However, many studies focus on the quality in biological process. Excessive changes in biological process would cause the consequence that is not expected. An example was shown in this study (Fig 1 A, B, C, D).

Recently, there has been growing interest in the research on quantitative stomata(Karavolias et al. 2023; Karavolias et al. 2024; Sakoda et al. 2020). Our study was also an initiating work in the research field where researchers study what effect of the quantitative stomata on plants morphology in genetical manipulation. In particular, it was strikingly that the lines with quantitatively increased stomatal density through genetic engineering indeed were able to possess the special outcomes. In this study, we used the 35S promoter to launch the overexpression of *FSTOMAGEN*, and chose the lines with different expression of *FSTOMAGEN*. This strategy has some limitations. Regarding the biotechnology, the representative widely known is the CRISPR-Cas9(Xing et al. 2014). It represents a more powerful tools for future research to directly edit genomes to modulate the differential expression of *FSTOMAGEN*.

The molecular mechanism of stomatal development has been well studied. Besides, stomata are convenient to observe and count, and many software tools for counting stomata has been developed in recent years(Fetter et al. 2019)(Wang et al. 2024). More crucially, reduced atmospheric CO_2_ and increased biomass could enable us to attach importance to the adjustment of stomata. Therefore, stomata could be a crucial model for the quantitative biology at the cell level, particularly in the field of plant science.

## Acknowledgement

Thanks for the cooperating of the experiments with Fu sang Liu. Thanks for the cooperating from the members in groups.

## Author contribution

Yong-yao Zhao designed the research in this article and wrote the article. Yong-yao Zhao leaded the experiment. Yong-yao Zhao analyzed the data. Yong-yao Zhao conducted the graphing and plotting.

---

Supplementary figure 1 The identification for the overexpression of FTOMAGEN A, The stomatal density for the different transgenic lines. B, The RT-PCR for the different transgenic lines. The photos and plotting are from(Zhao et al. 2022). The plant individuals used in this study are different from the individuals in the identification, but in the same line.

Supplementary figure 2 The photos for the different *Arabidopsis* lines with the *FSTOMAGEN* overexpression at the different date during the growth. The photos for the different *Arabidopsis* lines with different stomatal density at 19 day (A), 22 day (B). Stomatal density is shown above the photos.

Supplementary figure 3 The photos for the stomata in the leaf of different transgenic lines The scale bars in the photos were 100 μm.

Supplementary figure 4 The dry biomass for the different *Arabidopsis* transgenic lines under lower light intensity (300 μmol・m^-2^・s^-1^). A, The dry biomass of the different transgenic lines. B, The dry biomass of the three lines (14 line, 8 line and 9 line). C The A-ci curve of the two transgenic lines (14 line and 8 line).

Supplementary figure 5 The photosynthetic parameters for the three lines (14 line, 8 line and 30) A, A-q curve of the 14 line, 8 line and 30 line. B, Stomatal conductance of the three lines (14 line, 8 line and 30 line) at the different light intensity.

Supplementary table 1 The internal studentized residual and the external studentized residual for the relationship between leaf area and stomatal density.

Supplementary file The statistic test for the Figure 2 and 3.

